# Marsupial limb patterning redefines the necessity of lateral plate mesoderm subdivision for limb formation

**DOI:** 10.1101/2024.12.19.626501

**Authors:** Axel H Newton, Alexandra Leggatt, Ella R Farley, Karen E Sears, Aidan MC Couzens, Sara Ord, Andrew J Pask

**Affiliations:** The School of BioSciences, University of Melbourne, Victoria, Australia, 3000; Department of Ecology and Evolutionary Biology, University of California, Los Angeles, CA 90095; Colossal BioSciences, Dallas, Texas, USA

**Keywords:** LPM, heterochrony, heterotopy, somatic, splanchnic, marsupial

## Abstract

The tetrapod limb has long served as a model for elucidating molecular and cellular mechanisms driving tissue patterning, development and evolution. While significant advances have been made in understanding the drivers of limb initiation, outgrowth, and patterning, the early morphogenetic processes that transform the lateral plate mesoderm (LPM) into limb fields remain less resolved. Marsupial mammals provide a unique opportunity to investigate these foundational processes due to their accelerated forelimb development, driven by the functional demands of altricial neonates to crawl into the pouch at birth. Heterochronic formation of the forelimbs occurs prior to development of other surrounding structures, offering unparalleled insights into the plasticity of limb field specification. Here, we reveal that marsupial limb initiation and outgrowth bypasses physical subdivision of the LPM, a process previously considered critical for tetrapod limb formation. Instead, limb development proceeds through early activation of LPM-associated genes and proliferation before coelom formation, demonstrating remarkable morphogenetic plasticity. This evolutionary adaptation enables heterochronic limb development, redefining conserved processes to meet extreme functional constraints. These findings challenge previous models of tetrapod limb specification, highlighting the evolutionary plasticity of limb patterning mechanisms and reshaping our understanding of how selective pressures influence foundational developmental events.

## Introduction

The tetrapod limb—from wings to flippers and hands with opposable digits—has long been a model for understanding the molecular and cellular processes driving tissue patterning, development and evolution. Research in established models, such as the chicken and mouse, have clarified the molecular mechanisms driving limb development (for reviews, see ^1–3^), while comparative embryology across diverse non-model taxa has provided unique insights into how diverse limb types are patterned ^4–10^. Despite these advances, significant gaps remain in our understanding of the early morphogenetic events that transform the lateral plate mesoderm (LPM) into limb fields during early embryogenesis ^11,12^.

During early organogenesis, the limbs form via a series of morphogenetic events, arising from the LPM, a transient precursive tissue which underlies limb development ^13^. Initially, the mesodermal germ layer forms as bilateral sheets along the anteroposterior axis of the embryo, which then undergo mediolateral segmentation into paraxial, intermediate, and lateral (plate) domains ^14–17^. The LPM then undergoes further dorsoventral subdivision into somatic and splanchnic layers, which become separated by the coelomic cavity ^18,19^. This necessary transformation establishes physical division of the layers, allowing the somatic LPM to give rise to the body wall and limbs, while the splanchnic LPM forms the internal organs and viscera. Forelimb induction begins within the somatic LPM, via activation of the limb gene regulatory network through TBX5 ^20,21^. The limb-fated somatic LPM then undergoes epithelial-to-mesenchymal transition (EMT), producing the early limb mesenchyme ^22^. Spatial patterning and TBX5 activation in the somatic LPM-derived forelimb field is suggested through a complex interplay of genetic factors, including colinear expression of HOX genes ^23–26^, and secreted signalling pathways including WNT ^26–28^, retinoic acid (RA) ^29,30^ and BMP ^18,31^. However, signals from neighbouring structures, such as the developing neural tube, notochord, and somites, complicate the interpretation of individual signalling pathways during the critical window of limb induction. Studies across various vertebrate models have demonstrated that while some signals are necessary for limb induction, others are permissive, including RA signalling from the somites ^30,32,33^. The resulting complexity in signalling crosstalk has left ambiguity regarding the precise mechanisms that drive limb field specification and patterning.

Marsupial mammals offer a novel model system to disentangle the necessary molecular events underlying limb specification and patterning. Unlike eutherian mammals (including mouse and human), marsupials are born after a rapid gestation in a highly altricial state, yet require well-formed forelimbs to climb into the mothers pouch ^34,35^. To meet these functional demands, marsupials have evolved both a highly accelerated timeline of limb development (heterochrony), displaying accelerated expression of canonical limb genes (e.g., *TBX5, FGF10, FGF8, SHH*) ^36^; and anteroposterior expansion of their limb fields, allowing greater contribution of LPM-derived limb progenitors during limb outgrowth (heterotopy) ^37–40^. Notably, marsupial forelimb development occurs prior to the formation of surrounding structures, such as the somites and neural tube – ces of confounding signals in other vertebrate models ^40^ – thus developing in relative isolation. This unique developmental mode provides a rare opportunity to investigate the fundamental processes driving early limb field patterning. However, the early morphogenetic processes underlying limb field specification, LPM subdivision, and EMT of the somatic LPM have not been studied in marsupials, leaving unresolved questions about how accelerated limb development is achieved during its initial stages.

In this study, we use two distantly related marsupial models – the fat-tailed dunnart (*Sminthopsis crassicaudata*) ^41^ and gray short-tailed opossum (*Monodelphis domestica*) ^40^ – to dissect the cellular and molecular mechanisms underlying LPM formation and limb field specification in marsupial mammals. Through detailed visualization of key genetic markers of LPM specification and limb initiation, we investigate the cellular and molecular events underlying the heterochronic growth patterns in developing embryos. Our results highlight the isolated formation of the marsupial limb fields, where the forelimbs pattern and emerge prior to the formation of the neural tube, somites, and intermediate mesoderm. Furthermore, we show that forelimb outgrowth occurs independently of LPM subdivision, challenging existing models of LPM development. By studying these accelerated and divergent processes in marsupials, we can better understand the inductive signals driving the evolutionary plasticity of the tetrapod limb.

## Results

To characterise the onset of limb formation in marsupials, we compared the Australian marsupial *Sminthopsis crassicaudata* (fat-tailed dunnart) ^41^ and the American *Monodelphis domestica* (grey short-tailed opossum) ^42^, which last shared a common ancestor ∼80 million years ago. Embryos were assessed for the expression and localisation of four key genes marking early LPM specification and limb development ^43^: *PRRX1*, *FOXF1, TWIST1*, and *TBX5*. *PRRX1* is the first known marker of limb specification, activated in the undifferentiated LPM and persisting throughout the somatic LPM ^44–47^. *FOXF1* is similarly expressed in the undifferentiated LPM but subsequently becomes restricted to the splanchnic LPM ^18,48^. Following LPM subdivision, *TWIST1* is specifically activated in the somatic LPM just prior to *TBX5* expression ^43,49,50^, with *TBX5* serving as the definitive marker of forelimb initiation. Using whole-mount hybridisation chain reaction (HCR) and immunofluorescence, we determined that marsupial LPM specification and limb initiation occur rapidly over three developmental stages (stages 20–22, ∼8 hours), marking a distinct heterochronic shift compared to mouse and chicken development ^22,43^.

At stage 19, the trilaminar embryo is flat and tear-drop shaped, consisting of the three germ layers but only a few cells thick. It possesses a primitive streak and neural plate epithelium but lacks somites ^41^. Neither PRRX1 nor TBX5 could be detected at this stage (Figure S1), though a few hours later by stage 20 - where the embryonic disc starts to elongate posteriorly - the limb-fated LPM was present, marked by bilateral *PRRX1* and FOXF1 domains along the lateral aspects (Figure 1a,c). However, at this stage LPM subdivision had not yet occurred, as seen by largely TWIST1 negative limb fields, though few TWIST1 positive cells could be observed (Figure 1b) suggesting subdivision had begun. To determine whether the forelimb pathway had initiated, examination of *TBX5* expression revealed little to no overlap with *PRRX1* expression in the forelimb LPM of dunnart (Figure 1a), though *Monodelphis* showed feint overlapping *TBX5* with *PRRX1* in the limb fields (Figure 1c; also, where subdivision appeared to have occurred), suggesting that at this stage the limb pathway has just begun to be activated.

**Figure 1-.**
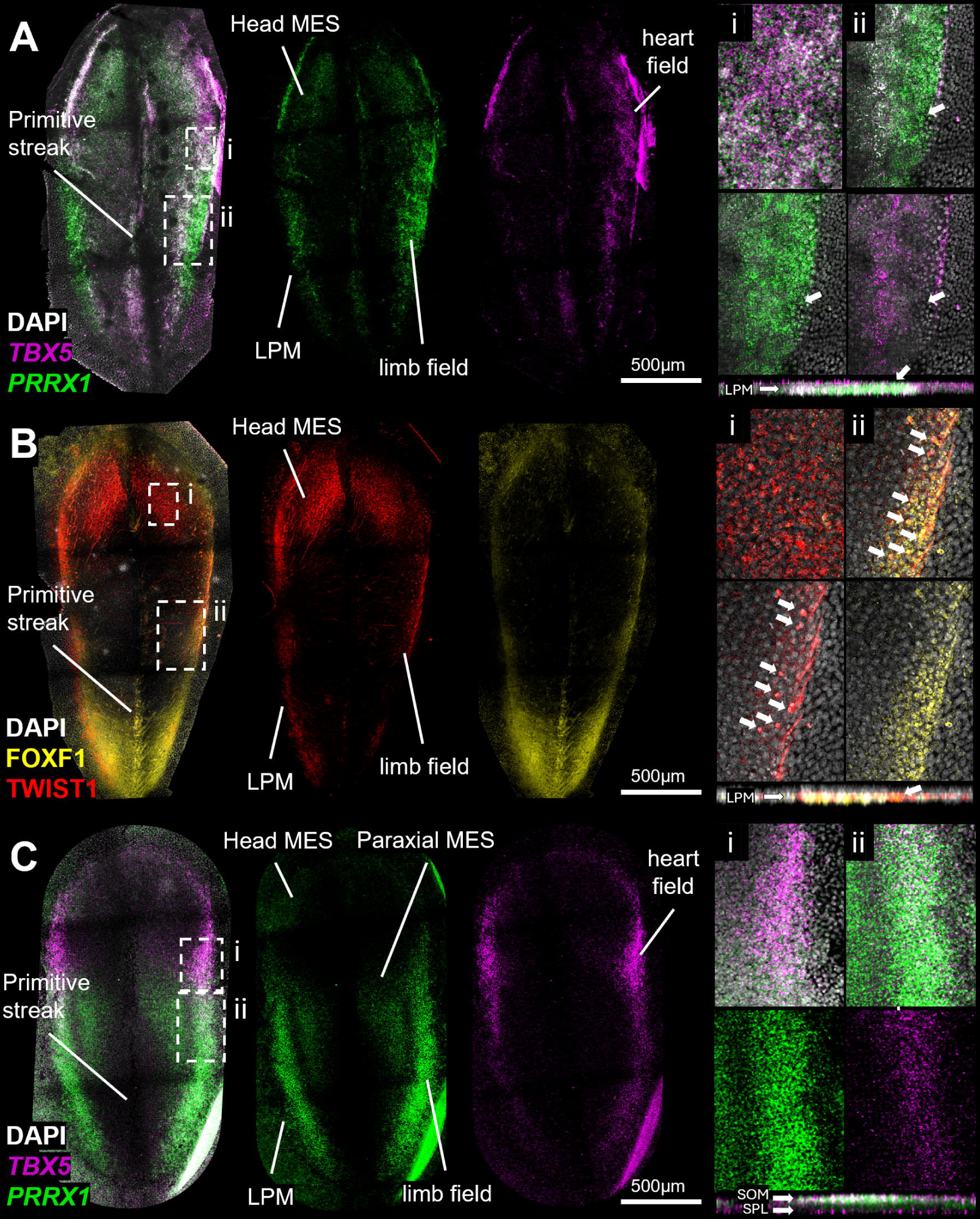
LPM specification in marsupial embryos. Stage 20 dunnart (A,B) and stage 21 opossum (C) primitive streak stage embryos at the time of LPM specification. The limb determining LPM has formed in these early embryos, marked by *PRRX1* and FOXF1. *TBX5* is expressed in the heart field (i) and feint expression can be seen in the putative limb field though this doesn’t particularly overlap with *PRRX1* (ii, white arrows), suggesting limb induction has not yet begun. B) FOXF1 positive cells are present throughout the LPM, and while strong TWIST1 is seen in the head mesoderm (i), only a subset of TWIST1 positive cells are present within the limb field (ii, white arrows), suggesting formation of the somatic LPM has just begun. C) Slightly older Monodelphis embryos display overlapping *PRRX1* and *TBX5* expression in the limb fields, as well as putative subdivision as seen through digital cross sections (ii, arrows), suggesting limb initiation has begun by this stage. LPM = lateral plate mesoderm; som = somatic LPM; spl = splanchnic LPM.

As the embryo progresses through stages 21 and 22, it elongates and forms distinct head folds despite the neural plate remaining flat, and the first somite pairs begin to condense within the paraxial mesoderm ^41^. At these stages, the bilateral forelimb fields can be observed along the sides of the embryo, marked by robust *PRRX1*, TWIST1 and *TBX5* positive cells demarcating the somatic LPM and early limb mesenchyme (Figure 2), though no hindlimb fields are yet present. Digital sections through the embryos revealed that the early forelimb fields were present as raised swellings, despite the rest of the embryo remaining flat and only a few cells thick (Figure 2). This raised questions about whether LPM subdivision had occurred, as this initial event typically causes dorsoventral expansion of the embryo through physical separation of the somatic and splanchnic LPM layers via formation of the coelomic cavity^18,22^.

**Figure 2-.**
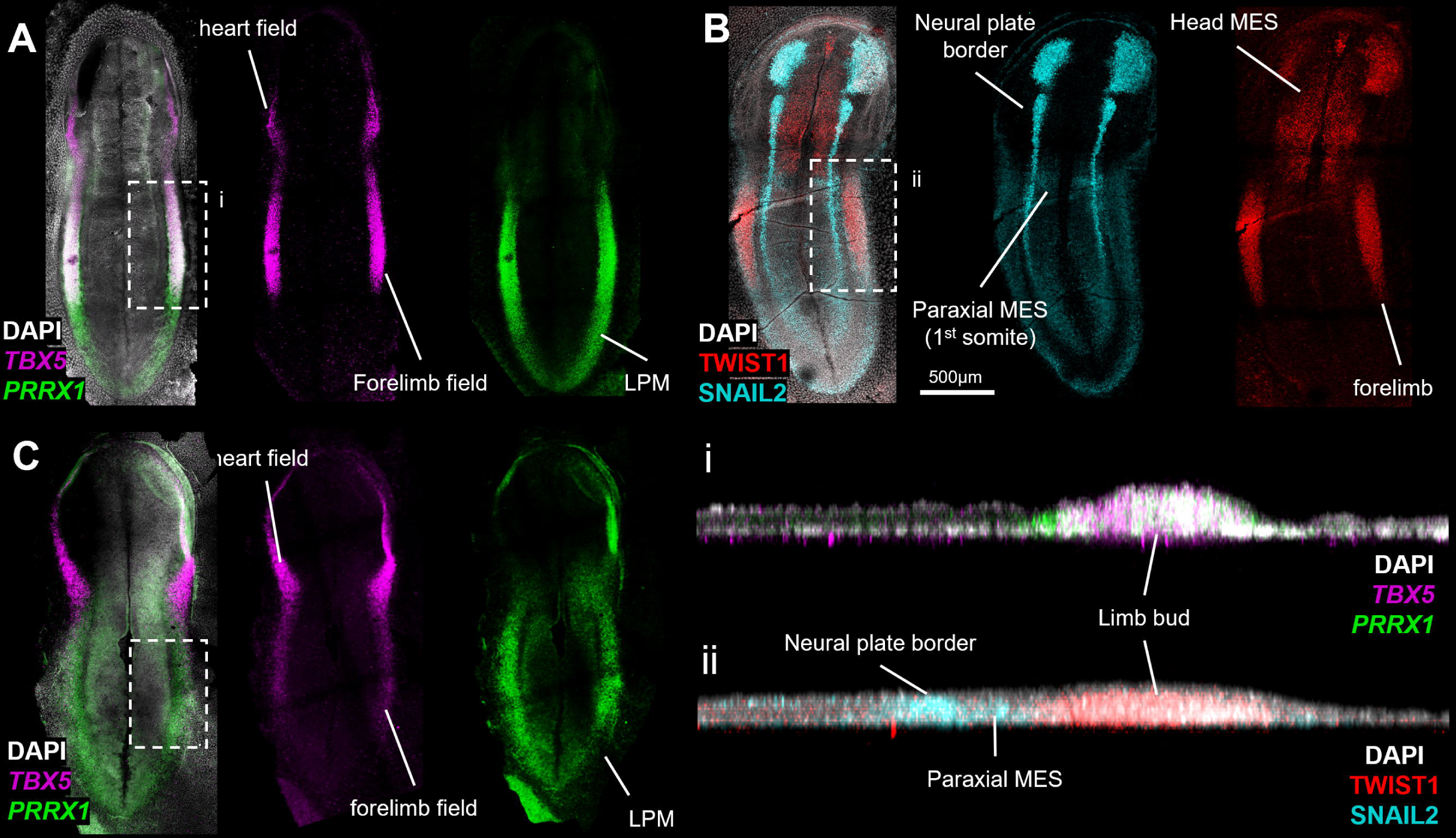
Limb induction in marsupial embryos. Stage 21/22 dunnart (A,B) and opossum (C) neurula stage embryos at the time of limb induction and outgrowth. The forelimb fields are rapidly specified into somatic LPM-derived limb buds, observed through co-localization of *PRRX1* and *TBX5* (A,C) and TWIST1 (B). The early limb buds can already be seen protruding from the flat embryo (digital sections i & ii). Notably, these thickened limb buds are present while the SNAI2 positive paraxial mesoderm remains rudimentary and is only just starting to condense into the first somite pair (B).

Histological analysis of ∼stage 20–22 forelimb fields revealed that marsupial limb formation occurs with superficial subdivision of the LPM without formation of the coelom. Remarkably, the LPM was observed to consist of a 1–2 cell-thick dorsal *TWIST1*-positive somatic layer attached to the underlying splanchnic layer (Figure 3a). In following stages, the lack of physical separation was clearly emphasized, remarkably showing proliferation and outgrowth of the limb bud mesenchyme (now 4-6 cells thick) while still attached to the FOXF1 and SNAI2 positive splanchnic LPM (Figure 3e,f). This process all occurred in the absence of somites, where the paraxial mesoderm could be observed as a thin layer of cells under the neural plate epithelium (Figure 3a,c,f,g). Examination of other cell or matrix markers, including E-cadherin, N-cadherin, Zo-1 and Laminin, suggested this superficial division was not accompanied by obvious EMT and cell state changes ^22^, nor formation of intermediate rosettes to physically divide the somatic and splanchnic layers ^19^ (Figure 3b-d,f-h). This demonstrated that during marsupial LPM formation, the somatic LPM becomes polarized and fated to form the limb mesenchyme, independent of physical subdivision typical of other vertebrates, notably mouse and chicken. Together, these findings from marsupial embryos challenge our existing understanding of the first stages of limb development. Here, physical subdivision of the LPM is not necessary for limb bud formation, the lack of somites at the time of limb outgrowth casts doubt on their role in limb initiation, and localized EMT of the somatic LPM does not appear to underlie the initial stages of limb bud outgrowth.

**Figure 3-.**
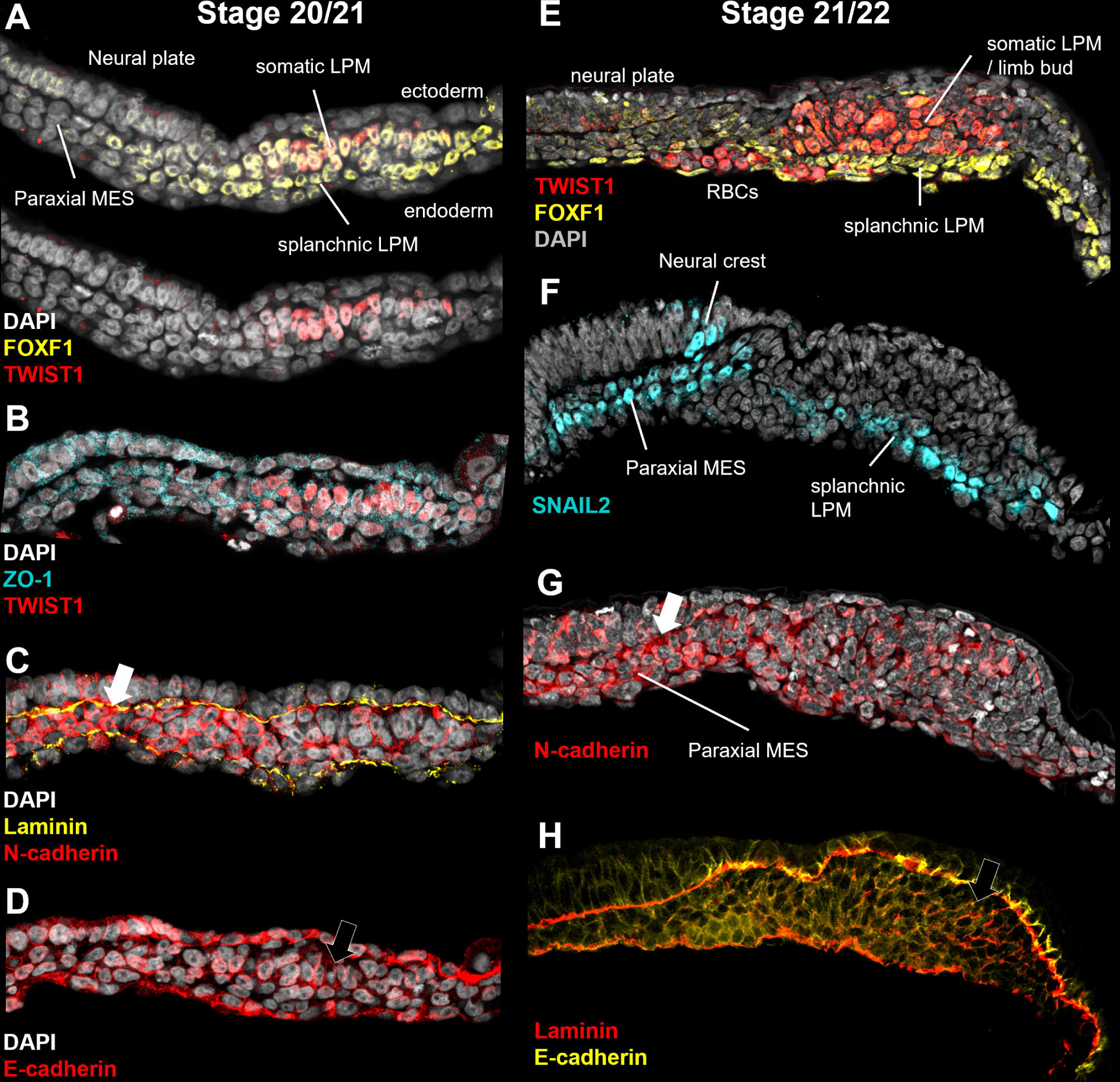
Subdivision of the lateral plate mesoderm occurs without physical separation. Tissue sections of the limb fields at the beginning of limb specification (stages 20/21, A-D) and during early outgrowth (stages 21/22, E-H), reveals subdivision of the marsupial LPM is incomplete without physical separation. At stage 20/21 the somatic LPM first forms, seen by TWIST1 localization, within the undifferentiated LPM, marked by co-localization with FOXF1 (A). These cells form as a small field still attached to the underlying splanchnic LPM, without clear apical markers (B), and retain an epithelial-like morphology marked by low N-cadherin, compared to the paraxial mesoderm (black arrow; C) and E-cadherin cell-cell junctions (black arrow; D). Following at stage 21/22, the TWIST1 positive somatic LPM has proliferated (E), though remains attached to the FOXF1/SNAI2 splanchnic LPM (F). These cells still possess low N-cadherin, compared with the paraxial mesoderm (white arrow), and do not break down the laminin boundary between the ectoderm, (G) and retain E-cadherin positive cell-cell junctions (black arrow) (H). LPM=lateral plate mesoderm; MES=mesoderm; RBCs = red blood cells.

The accelerated timetable of marsupial forelimb development became obvious during subsequent embryonic stages. By stage 24, both dunnart and opossum embryos possessed thickened forelimb buds with strong *TBX5* expression (Figure 4a,b), though these were comparative larger in the dunnart. Notably, physical subdivision of the LPM via separation by the coelom was now complete but occurring considerably after limb bud formation (Figure 4c). By this stage the rest of the embryo had started developing, notably the head, somites, tailbud, neural tube closure, and the developing heart tube, which was visible through the anterior *TBX5* field (Figure 4d). It should be noted that whilst somites have formed in the dunnart sample, these structures are still vague and poorly defined (Figure 4a). This lagging development of other key morphological structures emphasizes both the rapid formation of the forelimbs and how this developmental process occurs in physical isolation from other embryonic tissues during development.

**Figure 4-.**
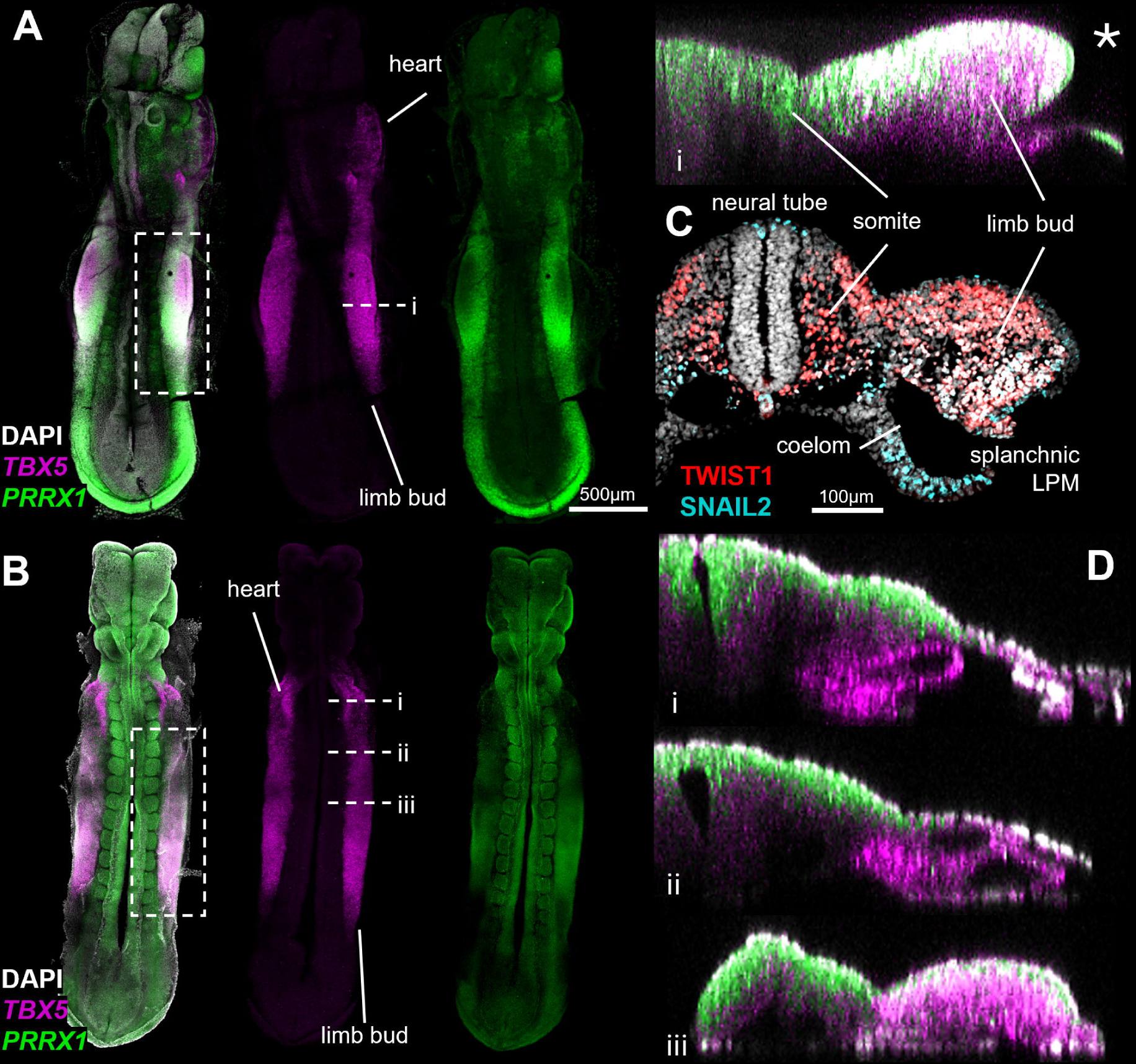
Outgrowth of the marsupial limb buds. Stage 24 dunnart (A) and opossum (B) embryos showing distinct outgrowth of the *PRRX1/TBX5* limb buds. At comparable stages, the dunnart limb buds are larger and more distinct than the opossum. C) At this stage, subdivision of the somatic (TWIST1) and splanchnic (SNAI2) LPM is complete, separated by the coelom, despite the limb buds being large and pronounced. D) Digital sections at anteroposterior levels of the *TBX5* domain in a *Monodelphis* embryos. This shows the progression from *TBX5* positive*; PRRX1* negative cardiac mesoderm, through to the outgrowing *TBX5*, *PRRX1* double positive limb bud.

To further contextualize this accelerated rate of development, known as heterochrony ^53^, we compared the marsupial patterns to the traditional chicken model. Marsupials undergo LPM specification and TWIST1-positive molecular subdivision of the somatic LPM whilst the embryo is a flat and featureless disc (Figure 1, 5a). In comparison, chicken subdivision precedes TWIST1 expression at a more developed stage ^43^, when the neural tube has closed and somites have condensed (Figure 5b). While less is known about the timing and anatomical context of subdivision in mice, it appears that this event precedes limb initiation with similar patterns to chicken ^48,54^ likely when the mouse has anywhere between 1 to 7 somites ^55^ (Table 1). Thus, marsupials exhibit extreme developmental heterochrony, initiating limb development in the absence of features such as a head, heart, closed neural tube, or fully formed somites at the limb level ^40^ (Table 1, Figure 5). On the other hand, at this same point of limb development, the chicken embryo is much more complex (22 somite stage; Figure 5), as are mice (8 somite stage), with the development of multiple other features such as the somites, the heart, and the head occurring concurrently with the development of the limb (Table 1). This is a striking difference to what is observed in marsupials, highlighting the extreme heterochronic acceleration of forelimb development amongst vertebrate species. The final stages of the initiation of limb development, the onset of limb proliferation and outgrowth, again occur at an earlier and less developed timepoint in marsupials when compared to mouse (21 to 29 somite stage) and chicken (26 to 28 somite stage). Overall, these comparisons help highlight and contextualise the level of heterochrony in marsupial species regarding the development of the forelimb.

**Figure 5-.**
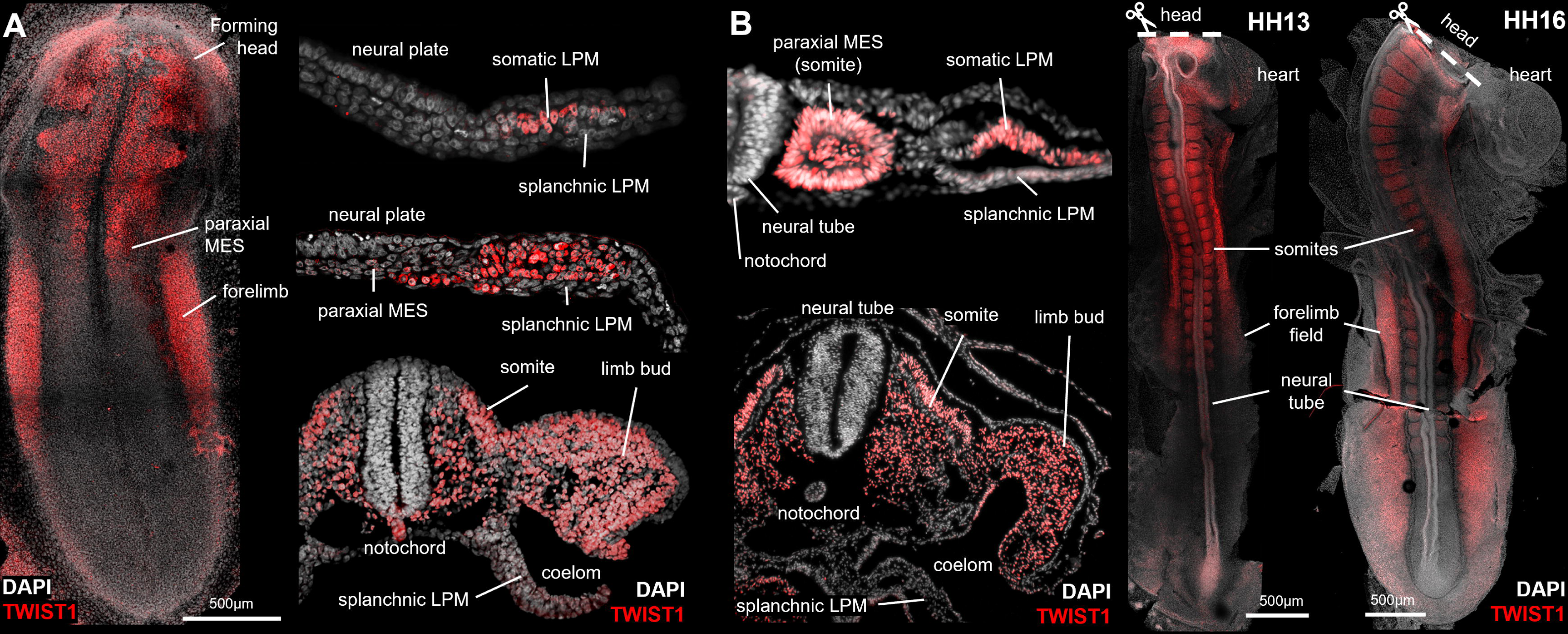
Heterochrony of LPM specification and limb development in marsupials. Comparisons of limb field specification in a stage 22 dunnart embryo (A) compared with similar stages in HH12 and 16 chicken embryos (B). Wholemount staining of TWIST1 shows marsupial limb fields form at highly accelerated stages, which lack other embryonic structures including the neural tube, somites, head and heart which are present in chicken at the time of limb field induction. Further, comparisons of LPM subdivision reveal a divergent mode of development in marsupials, where complete subdivision – present at the earliest stages of limb development in chicken – don’t occur until limb bud outgrowth in marsupials.

**Table 1.** Comparisons of the key features of embryos, matched according to stage of limb development. These four main features (head, neural tube, number of somites, and heart) were picked due to their well-characterised development, and their ease of visual comparison between stages and between species ^40,48,60,61^.

## Discussion

Owing to the functional requirements for the underdeveloped neonate to crawl into the mothers pouch, marsupials have evolved an accelerated timetable of forelimb development compared to other mammals, known as heterochrony ^34,35,40^. This heterochronic development is achieved not only through rapid outgrowth, but also enhanced patterning processes and allocation of limb progenitors, known as heterotopy. While this was previously attributed to early activation and expression of genes within the canonical limb pathway ^36,40^, we show that marsupials have evolved several atypical mechanisms to achieve their rapid forelimb development. This was first observed as formation of robust *PRRX1-* positive forelimb domains shortly after primitive streak formation, showing accelerated patterns compared to chicken and mouse^46,56^. Strikingly however, section histology through the early limb fields revealed that marsupial forelimb initiation and outgrowth occurs without physical subdivision or EMT of the somatic LPM ^18,19,22^, instead occurring through rapid proliferation of an attached, albeit molecularly distinct, somatic LPM layer. These findings challenge established views on how limb fields are produced in the early embryo, suggesting the mechanical processes underlying somatic LPM specification and limb development are more plastic than appreciated.

Subdivision of the LPM and formation of the coelomic cavity is a key morphogenetic event observed across vertebrates ^13^, and as far back as the chordate amphioxus ^57^. This process occurs along the anteroposterior as mediolateral wave, where (as observed in chicken) the lateral plate mesenchyme organizes into rosettes and epithelializes via a mesenchymal-to-epithelial transition (MET). This causes physical division of the somatic and splanchnic layers via embryonic coelom ^18,19^, allowing distinct lineage specification towards the limbs and viscera, respectively ^43^. During the first stages of limb outgrowth, the epithelial somatic LPM undergoes EMT and reverts into a proliferative mesenchyme (EMT), where this change in cell state facilitates extensive outgrowth of the limbs ^22^. However, our findings reveal that marsupials subvert these canonical processes. Instead, LPM subdivision occurs as a molecular polarization of dorsal somatic and ventral splanchnic layers which remain connected until later stages of limb bud outgrowth, without formation of intermediate rosettes ^19^. Moreover, the proliferating somatic LPM, or early limb buds, retain epithelial-like characteristics (notably E-cadherin positive cell-cell junctions) which no not display obvious evidence of EMT prior to, or during early limb outgrowth^22^. These observations challenge the long-held hypothesis that LPM subdivision, coelom formation, and somatic EMT are prerequisites for limb development. In the case of marsupials, the heterochronic shift in the timing of limb development driven by extreme functional constraints, appear to have redefined the morphogenetic boundaries of these conserved processes. This not only underscores the evolutionary plasticity of limb patterning, but also provides new insights into how these foundational events are regulated and adapted in response to distinct selective pressures.

The results of this study demonstrate that heterochronic formation of marsupial forelimb fields occur in effective isolation from other axial structures, which are present in other tetrapods at comparable stages. This notably includes the somites, which play roles in specifying mediolateral patterning of the mesoderm via BMP signalling ^14–17^, and are a source of secreted retinoic acid (RA) for *TBX5* activation in the somatic LPM ^30^. However, while RA is necessary in zebrafish ^32^ and chicken ^30^, it is dispensable in mice ^33^ making its precise role unclear. Though this was not examined in dunnart or opossum embryos, the absence of condensed somites during early limb formation supports the hypothesis that somitic RA is not universally required for mammalian limb development. This model paves the way for future analysis into other inductive signals thought to be necessary for forelimb induction, including collinear HOX genes ^23–26,30^, WNT ^26–28^ and BMP ^18,31^ signalling pathways – which have each been implicated in *TBX5* activation and limb induction ^30^. In *Monodelphis HOXB5* and *HOXC6* possess spatially increased domains correlating with their expanded limb field (spanning ∼8 somite pairs compared to ∼6 in mouse, human and chicken ^40^), implicating altered HOX expression in the enhanced formation of the limb domains, though its precise role in limb induction remains unclear. Through isolating the forelimb fields from the influence of other axial structures and focusing on conserved signalling pathways, marsupials offer a powerful system to disentangle the core processes driving limb field patterning and formation.

This study reveals novel insights into the early stages of limb patterning in marsupial mammals, and in turn challenges existing models of tetrapod limb development. We demonstrate that marsupial forelimb fields are specified in isolation through an accelerated and atypical sequence of events, including rapid specification, incomplete subdivision of the LPM and rapid outgrowth at the time of limb induction. These findings question the traditional view that LPM subdivision, coelom formation, and EMT are essential prerequisites for limb development. Moreover, the absence of somites and expanded forelimb field in marsupials downplay the role of retinoic acid (RA), while further implicating altered HOX expression as an early activator of the limb program. Together, these results show that the distinct trajectory of limb development in marsupials, shaped by evolutionary constraints, has produced plasticity in the fundamental processes underscoring limb development. Not only does this contribute to our broader understanding of the flexibility and diversity of developmental processes across tetrapods but emphasizes the value of using non-traditional models in developmental biology.

## Methods

### Dunnart husbandry and timed embryo collection

Dunnarts were obtained from an experimental colony run within the School of Biosciences. Female and male dunnarts were allocated into mating pairs in a 3:1 or 2:1 ratio of females to males. Detection of pregnancy for timed embryo collection was performed using recently described methods ^41^, where paired females were monitored for weight changes as the primary indicator of pregnancy, and females which showed consecutive days of weight increase after ovulation were used for embryo collection. Dunnarts were humanely killed by cervical dislocation, the paired uteri were then removed from the peritoneum and opened in DEPC-PBS. Pre-implantation embryos (<stage 28) were rolled out of the uterus, while implanted embryos were carefully cut out with their membranes intact. Embryos were transferred into 4% paraformaldehyde (PFA) and left to fix at 4°C overnight or over the weekend. Whole embryos fixed in PFA were washed in RNAse free 1X PBS with 0.1% Tween-20 (PBST) and then dehydrated in a 25%, 50%, 75%, and 100% methanol series (in PBS) on ice. Embryos were then stored in 100% methanol at −20°C until use.

### Wholemount and section fluorescent imaging

Gene expression analysis was performed using immunofluorescence and hybridization chain reaction (HCR) ^58^. Markers were chosen to represent specific stages in LPM and limb development, including PRRX1, SNAIL2, TWIST1, FOXF1, and TBX5.

Antibodies for transcription factors, cell and matrix proteins were prioritised, where available (Table 2). Methanol dehydrated embryos were transferred directly from 100% methanol into blocking buffer (1X PBS with 0.1% Tween-20 (PBST) with 3% BSA) and left to block for at least 1 hour, rotating at 4°C. Primary antibodies (Table 1) were diluted in PBST with 1% BSA and incubated with embryos overnight at 4°C while rotating. On the second day, embryos were washed with PBST to remove any unbound primary antibodies, then incubated with secondary antibodies (diluted at 1:500 in PBST with 1% BSA) at room temperature overnight in the dark while rotating. On the third day, embryos were washed in PBST and then incubated with DAPI (1:10,000 in PBST), before a final wash in PBST prior to mounting.

**Table 2.** Antibodies used in the study.

HCR probes (Molecular Instrument, Los Angeles, CA) were initially designed against target gene sequences from the dunnart (*Sminthopsis crassicaudata*) transcriptome ^59^. However, to ensure cross-reactivity between marsupials, probe sequences were BLASTed and retained only if they were specific to the gene of interest and had >85% similarity to *Monodelphis*. HCR was performed using the protocol provided by Molecular Instruments ^58^ with minor modifications based on the stage of the embryos. Briefly, embryos were rehydrated with the reverse methanol series (75%, 50%, 25%, 100% DEPC-PBST) on ice. Embryos between stages 24-25 were incubated with proteinase-K at room temperature for 5 minutes, then post-fixed in DEPC-PFA for 20 minutes at room temperature, while embryos at stages prior to stage 24 were not incubated with proteinase-K. All embryos were incubated with 10-30pmol of probes for *PRRX1* & *TBX5* in hybridisation buffer at 37°C overnight. On the second day, embryos were washed in wash buffer and DEPC-SSCT and were then incubated with 30pmol of H1 and H2 hairpins in amplification buffer at room temperature overnight. On the third day, embryos were washed in DEPC-SSCT and incubated with DAPI (1:10,000 in PBST), before being washed again in PBST.

### Mounting Embryos for imaging

Mounting was performed based on the size, stage and shape of the embryo. Mounting of early-stage embryos (stage 19-22) involved positioning the embryo flat on the slides with the dorsal surface facing upwards. Embryos often adopted the curvature of the vesicle and occasionally required further flattening through cutting the extra embryonic membranes. Embryos were then overlaid with glycerol mounting media (90% glycerol in 0.1 M PBS with 50 mg/ml propyl gallate) and a coverslip gently applied. Mounting of stage 23+ embryos required generation of custom silicon wells using aquarium-grade sealant. Embryos were positioned within the well and filled with the same fluorescent mounting media. A cover slip was then placed on top to seal the embryo within the chamber.

### Tissue Section Immunofluorescence

Whole embryos were transferred to 30% sucrose in PBS and allowed to sink, typically overnight. Embryos were then positioned within cryomolds in Tissue-Tek O.C.T. mounting media (ProSciTech) and stored at −80 until use. 15-20uM cryosections were cut and placed on alternating superfrost slides for successive immunostaining. Slides were washed in 1% Triton-X in 1X PBS (PBTX) and then blocked for at least 1 hour with 2% BSA in PBS at room temperature in a humidified chamber to prevent drying. Slides were then washed in PBS again and then incubated with primary antibodies diluted in PBS at 4°C overnight. On the second day, sections were washed in PBS and incubated with secondary antibodies (1:500 in PBS containing 1% BSA) at room temperature in the dark for 1 hour. Slides were then washed in PBS and incubated with DAPI (1:10,000 in PBS), before being washed again and mounted with ProLong Glass Antifade (Invitrogen) media and cover slipped.

### Imaging and analysis

Embryos and slides were imaged on a Nikon A1R confocal microscope with NIS-Elements software. Whole mount scans were captured with a 10x PL APO Lambda MRD00105 air objective (NA = 0.45), while tissue section images were imaged with a 40x PL FLUO MRH01401 oil objective (NA = 1.3). Due to the size of the wholemount samples, images were taken as large-image Z-stacks and were then stitched together in post-processing to make a single image. All image post-processing was performed using ImageJ (Fiji) for visualization, z-projection or z-slicing.

## Acknowledgements

The authors would like to thank The School of BioSciences (University of Melbourne) animal facility staff for the daily management of the Dunnart colony, particularly Shiralee Whitehead. We thank staff from the University of Melbourne Biological Imaging platform (BOMP) for microscopy usage and imaging assistance.

## Author contributions

AHN and AJP conceived the study. AL, ERF and AHN performed the experiments. AL and AHN reconstructed imaging data. AHN and AL analysed the data. ERF monitored dunnarts for pregnancy and collected embryos. AMC and KES generated and supplied opossum embryos. AHN, AL and AJP wrote the manuscript. All authors reviewed and gave final approval of the manuscript.

## Funding information

This research was conducted under research funding through Australian Research Council DP210102645 and DP160103683 to Andrew J. Pask, UoM ECR grant TP605149 to Axel H. Newton, generous philanthropic funding from the Wilson Family trust, and industry funding through Colossal BioSciences.

## Competing interests

Funding was provided by Colossal BioSciences (Texas, USA) for research costs and salaries related to the study.

## Data availability

All data is contained within the manuscript.

