## Supplementary figures and images for "Marsupial limb patterning redefines the necessity of lateral plate mesoderm subdivision for limb formation"

### Figure S1

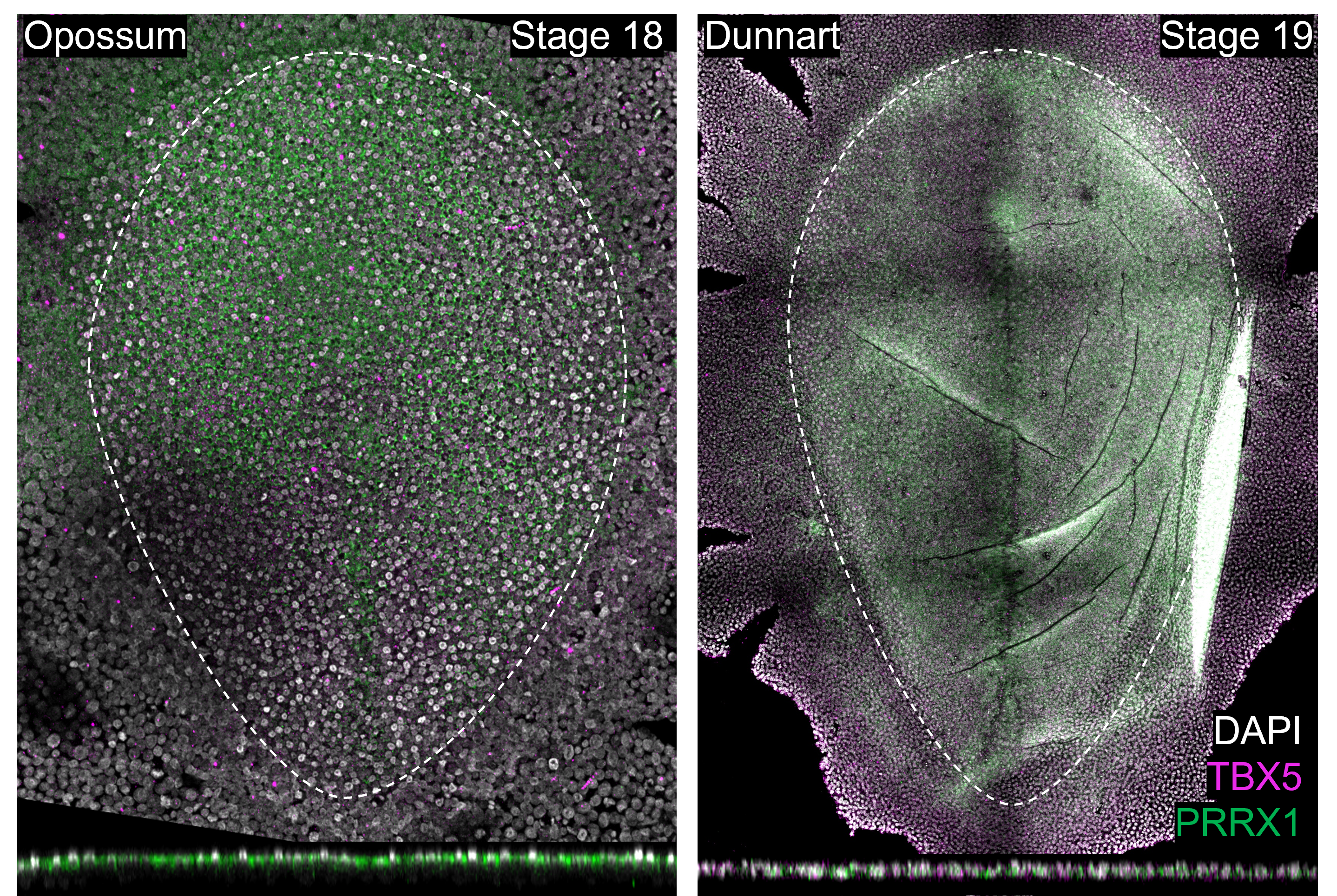
