## Supplementary material for "Marsupial limb patterning redefines the necessity of lateral plate mesoderm subdivision for limb formation": Table 1

| **Stage of limb development** | **Marsupial** | **Chicken** | **Mouse** |
| --- | --- | --- | --- |
| **Specification**  onset of *Prrxl* expression in the LPM | - Neural plate present (not yet folded into a tube) - Condensations of head mesoderm present - No somites - Vague heart field present - LPM not subdivided.  somatic and splanchnic layers not present | - Neural plate undergoing folding into the neural tube - Head is developed but not yet turned - 16 somite pairs present - Heart developing but not yet beating - LPM subdivided, somatic and splanchnic layers present | - NO data about Prrx1 expression in the LPM prior to specification of the somatic LPM and onset of Tbx5 expression is available. |
| **Initiation**   onset of Tbx5/TWISTl expression In the limb field | - Neural plate flat and still unfolded - First head fold bas become more developed - 1st somite pair present - Heart field more defined - LPM not subdivided, somatic and splanchlc layers present, but no coelom | - Neural tube closed - Head turned - 22 somite pairs present - Heart developed and beating - LPM subdivided and large coelom has formed | - Neural tube has begun close  Embryo is in process of turning, head is developing with branchial arches forming  8 somite pairs present  Heart is developed and beating  LPM subdivided and coelom present, but somatic and splanchnic layers only beginning specification |
| **Outgrowth**  onset of limb field proliferation | - Neural plate has thickened but still unfolded - Second head fold developed - 2nd somite pair present - Heart field still undeveloped, no heart visible - LPM subdivided, somatic and splanchnic layers present but no coelom present | - Neural tube closed - Head has turned and become more developed - 26 to 28 somite pairs present - Heart well developed and beating - LPM subdivided and large coelom present - Wing bud beginning to form | - Neural tube closed - Embryo bas completed turning - Head is well developed (forebrain closed and beginning separation of distinct vesicles) - 21 to 29 somite pairs present - Heart beating - LPM subdivided and coelom present. distinct somatic and splanchnic layers present |
